# The Hereditary and Epigenetic Fusion Gene Signatures of Multiple Myeloma

**DOI:** 10.1101/2022.12.05.519213

**Authors:** Ling Fei, Noah Zhuo, Degen Zhuo

## Abstract

Fusion transcripts are thought to be somatic and associated with cancer. However, they have been observed in healthy tissues at high recurrent frequencies. We have used SCIF (<u>S</u>plicing<u>C</u>odes <u>I</u>dentify <u>F</u>usion Transcripts) to analyze RNA-Seq data from 727 multiple myeloma (MM) patients of the MMRF CoMMpass Study. *MTG1*-*SCART1, SCART1*-*CYP2E1*, and *TPM4*-*KLF2* have been detected in 96.1%, 95.7%, and 92.2% of 727 MM patients and formed fusion gene signatures. *MTG1*-*SCART1* and *SCART1*-*CYP2E1* are read-through from the same locus and the two most recurrent epigenetic fusion genes (EFGs) out of 187 EFGs detected in ≥10% of 727 MM patients. *TPM4*-*KLF2* fusion gene, which was initially thought to be somatic, has been shown by a monozygotic twin genetic model to be a hereditary fusion gene (HFG) and the dominant genetic factor associated with MM. This work provides the first line of evidence that HFGs are the genetic factors and EFGs reflect the consequences between genetics and environments during development.

## Introduction

Multiple myeloma (MM) is a genetically complex and incurable blood malignancy characterized by the infiltration and growth of malignant plasma cells in the bone marrow^1^. Multiple myeloma affected approximately 427,000 people in 2013 and resulted in 79,000 deaths around the world^2^. In 2011, the Multiple Myeloma Research Foundation (MMRF) initiated the MMRF CoMMpass Study (MMRFCS), which performed comprehensive genomic analyses and followed each newly eligible patient longitudinally from initial diagnosis for ≥8 years^3^. Next-generation sequencing (NGS) of MM genomes had shown that MM was heterogeneous and was associated with significantly mutated genes and significant focal gene copy number gains and losses^4-6^. They resulted in definite, nonrandom chromosomal fusions of *IGHC* with the loci of *FGFR3, CCND1, CCND3, MAF*, and *MAFB* gene^7,8^. Analysis of 718 MMRFCS patients revealed that MM patients of European ancestries had higher mutations of *TP53* and *IRF4* than the African MM patients^9^. Cheynen et al. analyzed RNA-Seq data of a cohort of 255 newly diagnosed MM patients and identified 5.5 fusion genes, 29% of which were discovered in at least two patients^10^. Nasser et al. used RNA-Seq data from 806 samples with 704 matching WGS specimens and created 1192 of the observed fusions, a few of which were recurrent^11^. These two studies reflected overall obstacles and crises of RNA-Seq analyses because each software system discovered only small portions of fusion transcripts compromised by system errors and could not distinguish low expression fusion transcripts from “spurious” fusion transcripts^10,12-14^.

To overcome these problems and remove all system errors generated by experiments and software^15-17^, Zhou et al. used a structured table equal to 0.1% of human genome sequences. They implemented SCIF (<u>S</u>plicing<u>C</u>odes <u>I</u>dentify <u>F</u>usion Transcripts) to identify small subsets of fusion transcripts with maximum accuracy, eliminating “spurious” fusion transcripts^18^. Using SCIF, Zhou et al. discovered large numbers of fusion transcripts, among which the *KANSARL* (*KANSL1-ARL17A*) gene had been validated as the first predisposition (or hereditary) fusion gene specific to 28.9% of populations of European ancestry^18^. In addition, many studies have shown that fusion genes had been observed in cancer and healthy samples^8,19,20^, suggesting that fusion genes had many faces and required further investigation. To study fusion genes more precisely, we defined a hereditary fusion gene (HFG) as the fusion gene that offspring inherited from parents and excluded read-through fusion genes^21^ generated via transcriptional termination failure. Since environmental and physiological factors regulated read-through, we defined a read-through fusion gene as an epigenetic fusion gene (EFG)^22^. This report shows that *TPM4*-*KLF2, MTG1*-*SCART1*, and *SCART1*-*CYP2E1* genes have been detected in 92.2%, 96.1%, and 95.7% of 727 MM patients, respectively, and are the HFG and EFG signatures associated with MM. *TPM4*-*KLF2* is an HFG and *MTG1*-*SCART1* and *SCART1*-*CYP2E1* are EFGs.

## Materials and Methods

### Materials

#### Human multiple myeloma RNA-seq dataset

Multiple Myeloma Research Foundation (MMRF) initiated the MMRF CoMMpass Study in 2011. MMRF CoMMpass RNA-seq data (phs000748.v5.p4) were downloaded from controlled-access dbGaP (https://www.ncbi.nlm.nih.gov/gap). Seven hundred twenty-seven newly-diagnosed multiple myeloma patients were identified and used in this study. 99% of 727 RNA-seq data were from the bone marrow of multiple myeloma patients.

#### RNA-Seq dataset of Genotype-Tissue Expression (GTEx)

The RNA-Seq dataset of GTEx healthy blood samples had 427 healthy individual blood samples. RNA-Seq datasets of GTEx’s blood samples (dbGap-accession: phs000424.v7.p2) were downloaded from NCBI (https://www.ncbi.nlm.nih.gov/projects/gap/cgi-bin/study.cgi?study_id=phs000424.v7.p2).

#### Bone marrow samples

Four MM bone marrow samples and five healthy bone marrow samples were a gift from Prof. Yinxiong Li, Guangzhou Institutes of Biomedicine and Health, Chinese Academy of Sciences, China.

#### Primers

All primers used in this project were designed by Primer3Plus (https://www.bioinformatics.nl/cgi-bin/primer3plus/primer3plus.cgi).

#### Computers

All computations were performed on Linux Desktop computers with 8 to 12 GB memories.

## Methods

### Software

SCIF (<u>S</u>plicing<u>C</u>odes <u>I</u>dentify <u>F</u>usion Transcripts) had been described previously^18^.

### Recurrent Fusion Frequencies (RFF)

The recurrent fusion frequency of a fusion gene was equal to the number of samples with at least one isoform divided by the total number of samples.

### Classifications of fusion transcripts

The detailed protocols had been described previously^22^.

### Isolation of total RNA from bone marrows

Total RNAs of Bone marrow tissues were isolated by the Trizol reagent (Invitrogen, CA) described previously^18^.

### cDNA synthesis

cDNA synthesis was carried out as described by Zhou et al^. 18^.

### End-point PCR amplification

PCR amplification, DNA cloning, and sequencing were performed using the following primers: MS001F GAGACCATGGCTGACTACCT; MS001R GGACCAACACCACTCCGT; SC001F AATTCCACGAACATGAGCCC; SC001R CACATGAATGGGGCCAGAAG; TK001F GAGCGGCGCGAGAAAGTG; TK001R GTGCTTTCGGTAGTGGCG. The protocols and procedures had been described previously^18^. Prof. Yinxiong Li and his lab members performed RT-PCR and DNA sequencing.

## Results

### A brief review of enormous numbers of MMRFCS fusion transcripts

The MMRFCS RNA-Seq data (phs000748.v5.p4) consisted of 727 newly-diagnosed multiple myeloma (MM) patients. The MMRFCS RNA-Seq data had 144,069 million RNA-Seq reads, the average of which was 198.2 million RNA-Seq reads per patient. We used SCIF to analyze the MMRFCS RNA-Seq data. We identified 641,680 fusion transcripts supported by 1,405,768 unassembled raw RNA-Seq reads, each of which had 20-61 bp sequences identical to 5’ and 3’ genes plus splice site signals and correct orientations. Under these conditions, we performed extensive computational simulations and showed that our methods had removed potential experimental and computational artifacts. The false-positive rate was about 1% for cancer data and 1-2% for healthy samples. All false positives were from repetitive DNA sequences, pseudogenes, and alternative splicing.

We classified the fusion transcripts based on their recurrent frequencies observed in 727 MMRFCS MM patients to identify the potential fusion genes associated with MM. We discovered that 518 fusion transcripts ranged from 10% to 96.1%, twenty-one of which had ≥50% recurrent frequencies (Supplemental Table 1), suggesting that many fusion transcripts were highly recurrent and potential MM biomarkers. Based on their characteristics, fusion transcripts were initially classified into read-through fusion transcripts^21^ and fusion genes^23^ formed *via* genomic alterations, including deletions, inversions, intra-chromosomal, and inter-chromosomal translocations. These 518 fusion transcripts were classified into 284 fusion transcripts produced via genomic alternations and 234 read-through fusion transcripts. Their expression levels were higher than that of *KANSARL* isoform 1 and much lower than that of *KANSARL* isoform 2 (Supplemental Figure 1). The *IGHV-WHSC1* expression level was six-fold higher than that of *KANSARL* isoform 2 and was more than 100-fold higher than 99% of 518 fusion transcripts, suggesting that these 518 fusion transcripts were not highly expressed (Supplemental Figure 1). The average expression levels of the fusion genes were slightly lower than that of the read-through fusion genes. Out of 518 fusion transcripts, 284 fusion transcripts were encoded by 275 fusion genes generated by genomic alterations, which included deletions (5.1%), inter-chromosomal translations (69.1%), intra-chromosomal translocations (13.1%), and inversions (12.7%). Only four fusion genes had additional ≥10% recurrent isoforms, one of which was *NCOR2-UBC* having six additional isoforms.

On the other hand, 234 read-through fusion transcripts were encoded by 188 EFGs, 23.8% of which had additional isoforms with recurrent frequencies of ≥10%. The recurrent frequencies of the fusion genes produced by genomic rearrangements ranged from 10% to 92.2% of 727 MM patients, the average of which was 41.8%. In comparison, the EFGs ranged from 10% to 96.1% of 727 MM patients, the average of which was 52.9%. The former average recurrent frequency was 11.1% higher than the latter one. The higher EFG recurrent frequencies reflected that EFGs were more easily induced by cellular stresses, had a much longer time to be accumulated, and reflected many early events.

Since the *OAZ1-KLF2* fusion gene with a recurrent frequency of 58.1% was the second highest fusion gene generated by genomic alterations, the recurrent frequency of *TPM4*-*KLF2* was 34% higher than that of *OAZ1-KLF2*. Hence, *TPM4*-*KLF2* was the dominant fusion gene produced *via* genomic alterations. Based on their gene names, it was immediately apparent that *MTG1-SCART1* and *SCART1-CYP2E1* were two epigenetic (read-through) fusion genes (EFGs) from the same locus and were detected in 96.1% and 95.7% of 727 MM patients, respectively. They were at least 16.8% higher than the two closest *ACSS1*-*C20orf3* and *ERGIC1*-*RPL26L1* EFGs, with a recurrent frequency of 78.9% (Supplemental Table 1).

To evaluate if *TPM4*-*KLF2, MTG1*-*SCART*, and *SCART1*-*CYP2E1* were associated with MM, we used the GTEx blood samples as healthy control. Table 1 showed that *TPM4*-*KLF2, MTG1*-*SCART1*, and *SCART1*-*CYP2E1* were detected in 1%, 28.3%, and 10.8% of 427 GTEx blood samples, respectively. They were 92-fold, three-fold, and nine-fold higher than the GTEx’s ones. Statistical analysis showed these differences were significant (*p*<0.00002) and indicated that *TPM4*-*KLF2, MTG1*-*SCART1*, and *SCART1*-*CYP2E1* were associated with MM. These data confirmed that *TPM4*-*KLF2* was the dominant fusion gene formed *via* genomic deletion, while *MTG1-SCART1* and *SCART1-CYP2E1* were the dominant EFGs.

**Table 1.** Recurrent fusion frequencies of *TPM4*-*KLF2* HFG and *MTG1*-*SCART1* and *SCART1*-*CYP2E1* EFG in MM patients and GTEx blood samples.

| Fusion Gene ID | Multiple Myeloma (727) |  | GTEx Blood (427) |  | Folds |
| --- | --- | --- | --- | --- | --- |
|  | # | % | # | % |  |
| ***TPM4 -KLF2*** | 670 | 92.2 | 4 | 1 | 92** |
| ***MTG1 -SCART1*** | 699 | 96.1 | 121 | 28.3 | 3** |
| ***SCART1 -CYP2E1*** | 696 | 95.7 | 46 | 10.8 | 9** |
| Z-scores > 5.0; ** $p < 0.00002$ | | | | | |

### Validation of *TPM4*-*KLF2* fusion gene as a hereditary fusion gene (HFG) associated with MM

Fig.2a shows a schematic drawing of the standard genomic structure of *the TPM4*-*KLF2* region located on 19p13.1. A duplicated region underwent deletion and generated a putative *TPM4*-*KLF2* fusion gene genomic structure (Fig.1b), which was transcribed into putative *TPM4*-*KLF2* pre-mRNAs (Fig.1c). As shown in Fig.1d, *TPM4* exon 1b was spliced with *KLF2* exon 3 to produce the *TPM4*-*KLF2* main isoform. This *TPM4*-*KLF2* isoform was predicted to produce a putative 97 aa tropomyosin 4 and *kruppel*-like transcription factor 2 hybrid protein (Supplemental Figure 2). This hybrid protein had a head of 47 aa coiled-coil region of tropomyosin 4 and a tail of 50 aa region of *kruppel*-like transcription factor 2. The subcellular protein locations predicted by PSORTII^24^ were in the nucleus and less likely in mitochondria and were more similar to those of kruppel-like transcription factor 2 than tropomyosin 4’s ones.

**Figure 1:**
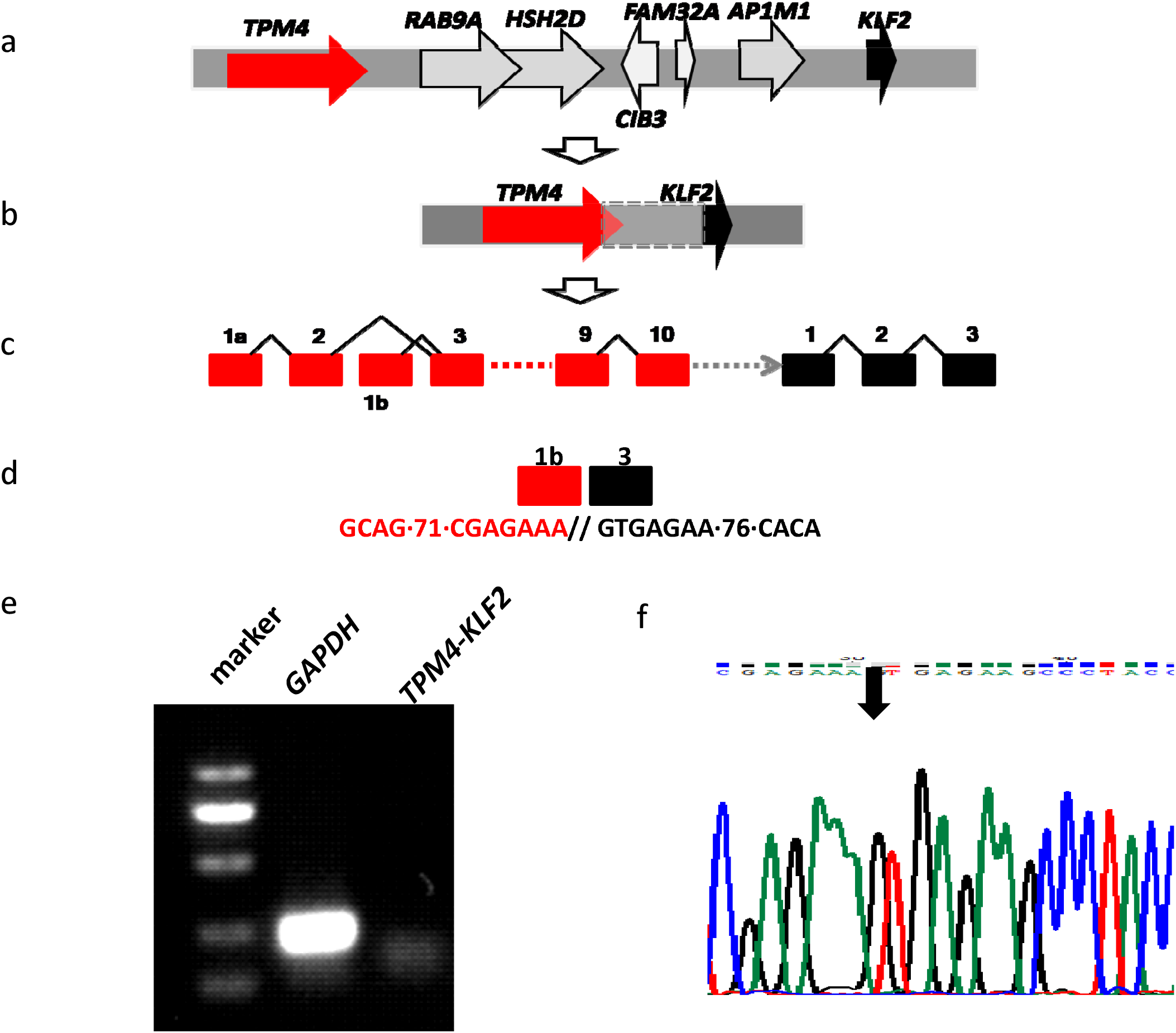
Validation of *TPM4-KLF2* fusion gene as an HFG. a). Schematic diagram of typical *TPM4*→*KLF2* genomic structures. Horizontal red and black arrows represented *TPM4* and *KLF2* genes, respectively. Horizontal gray arrows indicated genes between *TPM4* and *KLF2* genes; b). A potential deletion resulted in the *TPM4*-*KLF2* fusion gene; c) A putative *TPM4*-*KLF2* pre-mRNA structure. Black triangles represented introns. Numbers above the rectangles indicated exon positions of *TPM4* and *KLF2* genes, respectively; d). *TPM4*-*KLF2* mRNA was from their fusion junction. The sequences under the solid boxes were from RNA-Seq. e). RT-RNA PCR amplification of *TPM4*-*KLF2* fusion gene. f). DNA sequencing of cloned RT-PCR products validated the *TPM4*-*KLF2* fusion gene. Red and black boxes represented *TPM4* and *KLF2* exons, respectively. The red and black letters were *TPM4* and *KLF2* sequences, respectively.

**Figure 2:**
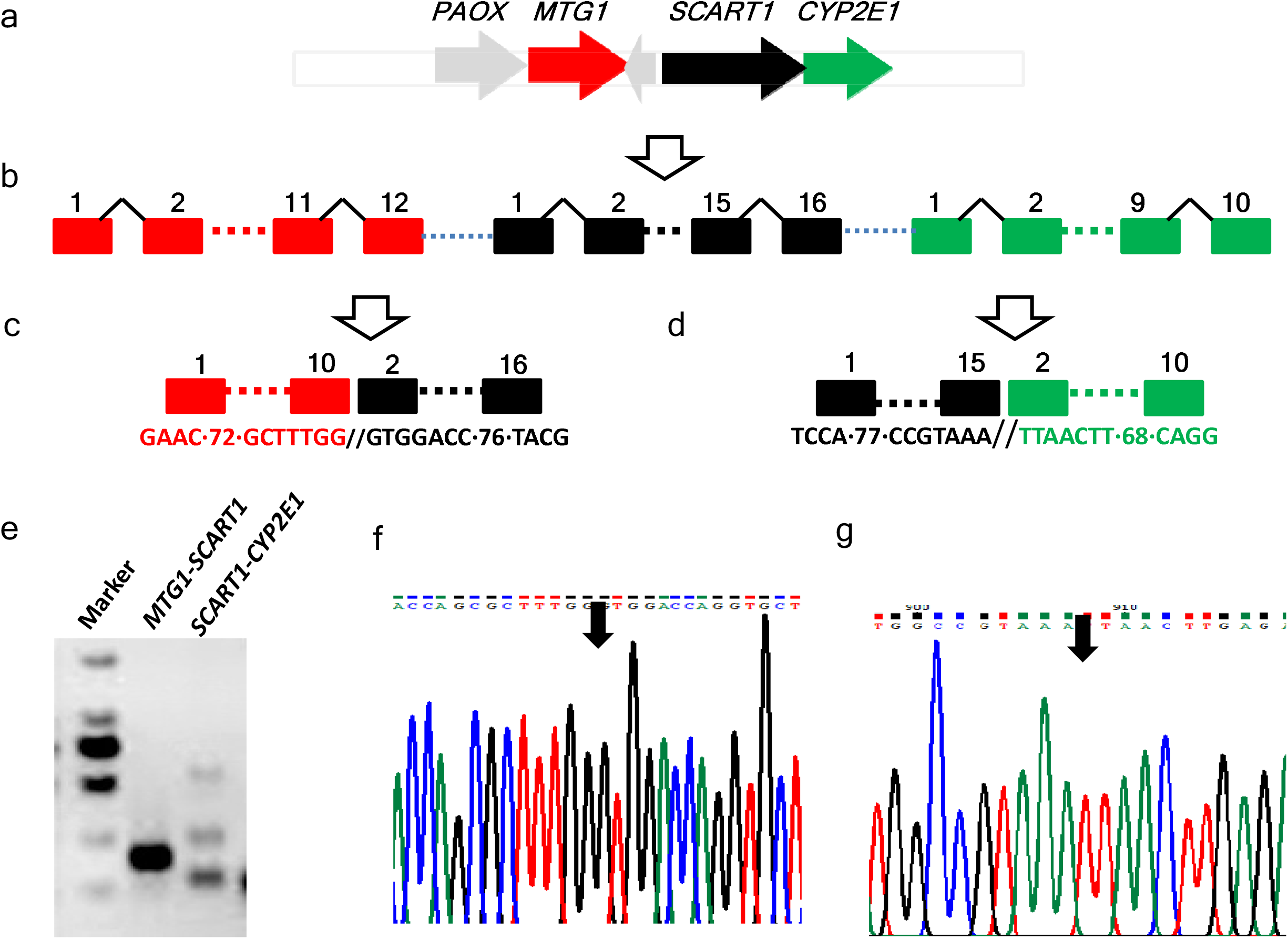
Characterizations of *MTG1-SCART1* and *SCART1-CYP2E1* EFGs. a) Schematic diagram of the genomic structure of PAOX→*MTG1*→*SCART1*→*CYP2E1*. Horizontal gray, red, black, and green arrows represented *PAOX, MTG1, SCART1,* and *CYP2E1*, respectively; b) Failed transcriptional terminations may produce *MTG1*→ *SCART1*→*CYP2E1* pre-mRNA transcripts. Red, black and green rectangles represented *MTG1, SCART1*, and *CYP2E1* exons. Black triangles represented introns; c) *MTG1*-*SCART1* fusion transcripts. Red and black rectangles indicated *MTG1* and *SCART1* exons, respectively. The red and black letters were *MTG1* and *SCART1* exon sequences, respectively. The sequences under the rectangles were from RNA-Seq; d) *SCART1-CYP2E1* fusion mRNA. The sequences under the rectangles were from RNA-Seq. Black and green rectangles indicated *SCART1* and *CYP2E1* exons, respectively. The black and green letters were *SCART1* and *CYP2E1* exon sequences, respectively; e) RT-PCR amplification of *MTG1*-*SCART1* and *SCART1-CYP2E1* fusion transcripts of MM patients; f). DNA sequencing of cloned RT-PCR products validated the *MTG1*-*SCART1* fusion gene. g). DNA sequencing of cloned RT-PCR products validated the *SCART1-CYP2E1* fusion gene. Vertical arrows indicated the fusion junctions.

As Table 1 showed, the *TPM4-KLF2* fusion gene had been observed only in four individuals out of 427 GTEx blood samples, which was about 1% of the GTEx’s blood samples, suggesting that *TPM4*-*KLF2* fusion transcripts were produced from a potentially rare somatic deletion. To validate the *TPM4*-*KLF2*, we first selected four MM patients as a starting point. We independently performed RT-PCR amplification on the bone marrow RNAs isolated from the four MM patients, all of whom were *TPM4*-*KLF2* positive. Fig.1e showed one of the four RT-PCR amplifications. All four RT-PCR products were cloned and sequenced independently. Fig.1f showed one of four *TPM4*-*KLF2* sequences and demonstrated that the sequence had the identical *TPM4*-*KLF2* fusion junction observed from RNA-Seq data. To study *TPM4*-*KLF2* fusion gene frequencies in healthy control, we performed RT-PCR analyses on the five healthy bone marrow samples, and consequently, five out of the five healthy controls were shown to be *TPM4*-*KLF2* positive. The healthy samples being *TPM4*-*KLF2* positive were significantly higher than the GTEx one. A literature review showed that *TPM4-KLF2* was first reported in acute lymphoblastic leukemia by Roberts et al.^25^. Locher et al. reported that *TPM4*-*KLF2* was detected in 30% of acute myeloid leukemia samples, and all three normal bone marrow samples^26^.

These high healthy *TPM4*-*KLF2* frequencies contradicted our original idea that *TPM4*-*KLF2* was a somatic genomic alteration and led us to speculate that *TPM4*-*KLF2* was similar to the hereditary *KANSARL* fusion gene. Such high *TPM4-KLF2* frequency compelled us to use monozygotic (MZ) twins with identical genetic materials as a genetic model to study human hereditary fusion genes (HFGs) systematically ^22^. We had identified a total of 1180 HFG from 37 pairs of MZ twins^22^. Interestingly, *TPM4*-*KLF2* was detected in 15 (40.5%) of 37 pairs of MZ twins and 54.1% of 74 twin siblings^22^. These data had mathematically ruled out that *TPM4*-*KLF2* was from a somatic genomic deletion. Hence, we could determine that *TPM4*-*KLF2* was a hereditary fusion gene.

To investigate how many fusion genes generated via genomic alterations were potential HFGs, we first overlapped the top five fusion genes with recurrent frequencies of ≥50% with 1180 MZ twin HFGs. Four of them were present in the MZ twin HFGs and were *TPM4*-*KLF2, OAZ1*-*KLF2, TRNAN35*-*FAM91A3P*, and *FOSB*-*KLF2*. Only *VPS37B-UBC* was not present among 1180 MZ HFGs. Next, we overlapped 275 fusion genes with recurrent frequencies of ≥10% with 1180 MZ HFGs, respectively. 87 (31.6%) of 275 fusion genes overlapped with the MZ HFGs and hence, were HFGs (Supplemental Table 2). Since the MZ HFGs were only tiny portions of potential 5×10^9^ HFGs^22^, real HFGs associated with MM would be significantly higher. Since somatic fusion gene *IGHV-WHSC1* had only 8.5% recurrent frequency and four of the five fusion genes with recurrent frequencies of ≥50% were HFGs, we estimated that ≥80% of 275 fusion genes generated via genomic alterations were inherited from their parents and not from somatic genomic abnormalities, suggesting that there were ≥220 potential HFGs associated with MM. These large numbers of HFGs and potential HFGs associated with MM suggested that multiple myeloma was a complex genetic disease.

### Characterization of *MTG1-SCART1* and *SCART1-CYP2E1* EFGs

The most frequently observed fusion genes were *MTG1*-*SCART1* and *SCART1*-*CYP2E1*, detected in 96.1% and 95.7% of 727 MMRFCS MM patients. Based on their gene names, it was immediately apparent that *MTG1*-*SCART1* and *SCART1*-*CYP2E1* fusion transcripts were read-through products from the same locus. Fig.2a showed that *MTG1, SCART1*, and *CYP2E1* genes (red, black, and green rectangles represented *MTG1, SCART1*, and *CYP2E1*) clustered together located on the 10q24.3 plus-strand. *MTG1, SCART1*, and *CYP2E1* genes encoded mitochondrial GTPase 1, scavenger receptor 1, and cytochrome P450 2E1. Failed transcriptional terminations produced a single read-through pre-mRNA containing three genes or two separate *MTG1*-*SCART1* and *SCART1*-*CYP2E1* read-through pre-mRNAs (Fig.2b).

Fig.2c showed that *cis*-splicing of *MTG1*-*SCART1* read-through pre-mRNAs resulted in *MTG1*-*SCART1* isoform (designed as *MTG1*-*SCART1* isoform 1) as discovered from the above RNA-Seq analysis. *MTG1*-*SCART1* isoform 1 would contain the first ten *MTG1* exons and the last 14 *SCART1* exons and produce an open frame of 461 aa hybrid polypeptide, which consisted of 256 aa mitochondrial GTPase 1 and 205 aa scavenger receptor 1 with an isoelectric point of 10.17 and molecular weight of 50 kd. PSORTII^24^ predicted 34.8%, 30.4%, and 17.4% in the cytoplasm, nucleus, and mitochondria, respectively, significantly different from mitochondrial GTPase 1 of 73.9% and 13% in the cytoplasm and nucleus^21^. On the other hand, Fig.2d showed that the exon 1-15 of *SCART1* was putatively spliced together with the exon 2-10 of *CYP2E1* to produce *the SCART1-CYP2E1* isoform (named *SCART1*-*CYP2E1* isoform 1). Since the *SCART1*-*CYP2E1* fusion junction sequences were located at the *SCART1* 3’ UTR region (Fig.2d), the *SCART1*-*CYP2E1* isoform 1 would produce a normal scavenger receptor 1. Based on their fusion junctions, *MTG1*-*SCART1* and *SCART1*-*CYP2E1* were predicted to produce a mitochondrial GTPase 1-scavenger receptor 1 hybrid protein and a normal scavenger receptor 1, respectively.

To validate *MTG1*-*SCART1* isoform 1 and *SCART1*-*CYP2E1* isoform 1, we independently isolated total RNAs from the bone marrow of four MM patients. Fig.2e showed that one of four MM patient samples was used for RT-PCR amplification on *MTG1*-*SCART1* and *SCART1*-*CYP2E1*. Four MM patients’ RT-PCR products were isolated, cloned, and sequenced independently. Fig.2f showed that one of four cloned RT-PCR *MTG1*-*SCART1* sequences had identical fusion junction sequences from RNA-Seq. Similarly, Fig.2g showed that one of four MM patients had fusion sequences identical to *SCART1*-*CYP2E1* isoform 1.

Table 1 showed that *MTG1*-*SCART1* and *SCART1*-*CYP2E1* in MMRFCS MM patients in the GTEx blood samples were three-folds and nine-folds more frequent than those in the GTEx blood samples. *MTG1*-*SCART1* and *SCART1*-*CYP2E1* were detected in 28.3% and 10.8% of the GTEx blood samples. The former was much more frequently detected than the latter and 2.6 folds higher, suggesting that *MTG1*-*SCART1* and *SCART1*-*CYP2E1* were not co-expressed and differentially regulated in the healthy population. Since *TPM4*-*KLF2, MTG1*-*SCART1*, and *SCART1*-*CYP2E1* in MMRFCS MM patients had almost identical recurrent frequencies, *TPM4*-*KLF2* might have affected the expression of *MTG1*-*SCART1* and *SCART1*-*CYP2E1*. To investigate if *TPM4*-*KLF2* was associated with their expression, we compared their gene expression patterns with those of 74 MZ twin siblings. Supplemental Table 3 showed that *MTG1*-*SCART1* and *SCART1*-*CYP2E1* were detected in 48.6% and 18.9% of 74 MZ siblings, respectively. The *MTG1*-*SCART1* recurrent frequency was 2.6 folds of that of *SCART1*-*CYP2E1*.

In comparison, *MTG1*-*SCART1* and *SCART1*-*CYP2E1*, detected in 96.1% and 95.7% of the MM patients, had almost identical recurrent frequencies and were almost co-expressed. The differences between MM patients and MZ twins suggested that their expressions were not directly associated with *TPM4*-*KLF2*. Other genetic and environmental factors contributed to their expression. Therefore, *MTG1*-*SCART1* and *SCART1*-*CYP2E1* were the unique EFG signatures specific to MM and could be used as potential biomarkers to monitor MM development and progression.

Like a traditional gene, these EFGs could undergo alternatively splicing. Supplemental Table 4 showed that *MTG1*-*SCART1* encoded an additional 17 *MTG1*-*SCART1* isoforms, while *SCART1*-*CYP2E1* had additional eight isoforms (Supplemental Table 5). In addition, we found that *MTG1-CYP2E1* fusion transcripts had been detected in 204 (28.1%) of 727 MM patients (Supplementary Table 1). However, no single copy of the *MTG1-CYP2E1* fusion transcript had been observed in the GTEx blood samples, suggesting that *MTG1-CYP2E1* reflected the MM progression. In a recent report, *MTG1-CYP2E1* was a potential pathogenetic link between angiomyofibroblastoma and superficial myofibroblastoma in the female lower genital tract^27^. Since *PAOX*-*MTG1* had been annotated as a *PAOX*-*MTG1* EFG^21^, the *PAOX*-*CYP2E1* locus had produced four EFGs, which underwent extensive alternative splicing and dramatically increased biological and functional diversities.

## Discussion

This report used SCIF to analyze 727 MMRFCS RNA-Seq data and identified 641,680 fusion transcripts. Based on their recurrent frequencies, we discovered 518 fusion transcripts ranging from 10% to 96.1%, suggesting that enormous fusion genes may be associated with MM. Based on the types of fusion transcripts, fusion genes were classified into read-through^21^ and fusion genes generated via genomic alterations^14^. The expression levels of both EFGs and fusion genes were between those of *KANSARL* (*KANSL1-ARL17A*) isoform 1 and 2^18^, suggesting that these recurrent EFGs and fusion genes were not highly expressed. It seemed that they had undergone a natural selection and were tamed. Among the high recurrent fusion genes, *TPM4*-*KLF2, MTG1*-*SCART*, and *SCART1*-*CYP2E1* fusion genes were detected in ≥92% of 727 MMRFCS MM patients, had much higher recurrent frequencies than the rest of them, and formed the fusion gene signatures associated with MM.

When we performed RT-PCR amplifications on the healthy control, all five individuals were also *TPM4*-*KLF2* positive. This high *TPM4*-*KLF2* frequency led us to speculate that the *TPM4*-*KLF2* fusion gene was similar to the *KANSARL* predisposition fusion gene^18^ and compelled us to use MZ twins to study HFGs systematically. We analyzed RNA-Seq data of 37 pairs of MZ twins and discovered 1180 HFGs^22^. Coincidently, one of many HFGs associated with MZ twin inheritance was *TPM4*-*KLF2*, which was detected in 15 of 37 pairs of MZ twins and 54.1% of 74 twin siblings^22^. The MZ twins’ high *TPM4*-*KLF2* recurrent frequency had mathematically ruled out that *TPM4*-*KLF2* was a somatic fusion gene. Hence, *TPM4*-*KLF2* was determined to be a hereditary fusion gene. *TPM4*-*KLF2* was first reported in 2012^25^ and was later found in 30% of acute myeloid leukemia (AML) samples and all three normal bone marrow samples^26^, suggesting that *TPM4*-*KLF2* was associated with AML. Hence, *TPM4*-*KLF2* was a broad-spectrum HFG associated with multiple types of cancer and MZ genetics. *PIM3-SCO2*^*26,28*^, *NCOR2-UBC*^*26*^, and *OAZ1-KLF2*^*25,26*^ were previously-studied cancer fusion genes, also associated with the MZ twin inheritance and HFGs^22^. Hence, there was no such thing as a good HFG or a bad HFG whether an HFG caused what type of disease depended on other HFGs, developments, and environments.

To further understand the HFG role in MM, we showed that *TPM4*-*KLF2, OAZ1*-*KLF2, TRNAN35*-*FAM91A3P*, and *FOSB*-*KLF2* were HFGs. *TPM4*-*KLF2, OAZ1*-*KLF2*^*25,26*^, and *FOSB*-*KLF2* with recurrent frequencies of≥50% suggested that disrupting *KLF2* expression may be associated with MM. To check if there were more HFGs associated with MM, we overlapped 275 fusion genes of ≥10% were overlapped with 1180 MZ HFGs. 87 (31.6%) of them were HFGs. Since four of the top five fusion genes were HFGs and *IGHV-WHSC1* had only 8.5% recurrent frequency, ≥80% of 275 fusion genes were predicted to be putative HFGs. Many previously-characterized somatic fusion genes, such as *IGHV-WHSC1*^*8*^ and driver oncogenes, had much lower recurrent frequencies and were much later events. Hence, enormous numbers of HFGs were involved in MM and suggested that MM was a complex genetic disease, and *TPM4-KLF2* HFG was the dominant genetic factor identified so far and a potential therapeutic target.

*MTG1*-*SCART1* and *SCART1*-*CYP2E1* were two EFGs from the same locus detected in 96.1% and 95.7% of MM patients, suggesting that both were almost co-expressed in MM. The *MTG1*-*SCART1* and *SCART1*-*CYP2E1* main isoforms potentially encoded a novel mitochondrial GTPase 1-scavenger receptor 1 hybrid protein and a normal scavenger receptor 1 protein, respectively, suggesting that both main isoforms had two different potential consequences. In addition, *MTG1*-*SCART1* and *SCART1*-*CYP2E1* EFGs had been found to have 18 and 9 isoforms detected in MM, respectively, which dramatically increased potential biological diversities. We compared data from MM patients and MZ twins to study if *TPM4*-*KLF2* was directly associated with their expression. Supplemental Table 3 showed that *MTG1*-*SCART1* and *SCART1*-*CYP2E1* were detected in 48.6% and 18.9% of 74 MZ twin siblings. The former recurrent frequency was 2.6 folds of the latter, indicating that their patterns in MZ twins significantly differed from those in the MM patients. *MTG1*-*SCART1* and *SCART1*-*CYP2E1* had almost identical recurrent frequencies in MMRFCS MM and were the unique EFG signature specific to MM.

*TPM4*-*KLF2* HFG and *MTG1*-*SCART1* and *SCART1*-*CYP2E1* EFGs had formed the HFG and EFG signatures of MMRFCS MM. In order to initiate MM, cells must have *TPM4*-*KLF2* HFG and a set of other MM HFGs (Fig.3a). Expression of these sets of HFGs resulted in read-through pre-mRNAs to produce EFGs such as *MTG1*-*SCART1* and *SCART1*-*CYP2E1* EFGs and maybe activated some dormant HFGs (Fig.3b). Without treatments and interventions, these cells expressed excessive EFGs and HFGs, resulted in genomic alternations, and produced somatic fusion genes. Cells underwent natural selection to lead to MM developments and progression (Fig3c). In comparison, if healthy individuals had only *TPM4*-*KLF2* HFG and no other MM HFGs (Fig.3d), cells expressed much fewer EFGs and could tolerate more stresses (Fig.3e). Hence, they maintained typical genomic structures and produced few somatic gene mutations (Fig.3f). Fig.3 showed that individuals must have both a set of MM HFGs and favorable environments to initiate MM. Without any one of them, the cells would be divided typically. Therefore, we could use a drop of blood or a few cells to screen HFGs and EFGs associated with MM as early as the middle of pregnancy to monitor MM development and progression. Meanwhile, we could have enough time to develop different approaches or therapeutic drugs to prevent cell progression to MM. Hence, applications of HFGs and EFGs will potentially revolutionize cancer research, diagnosis, and treatments and potentially mark the beginning of a new era of cancer and other disease research.

**Figure 3.**
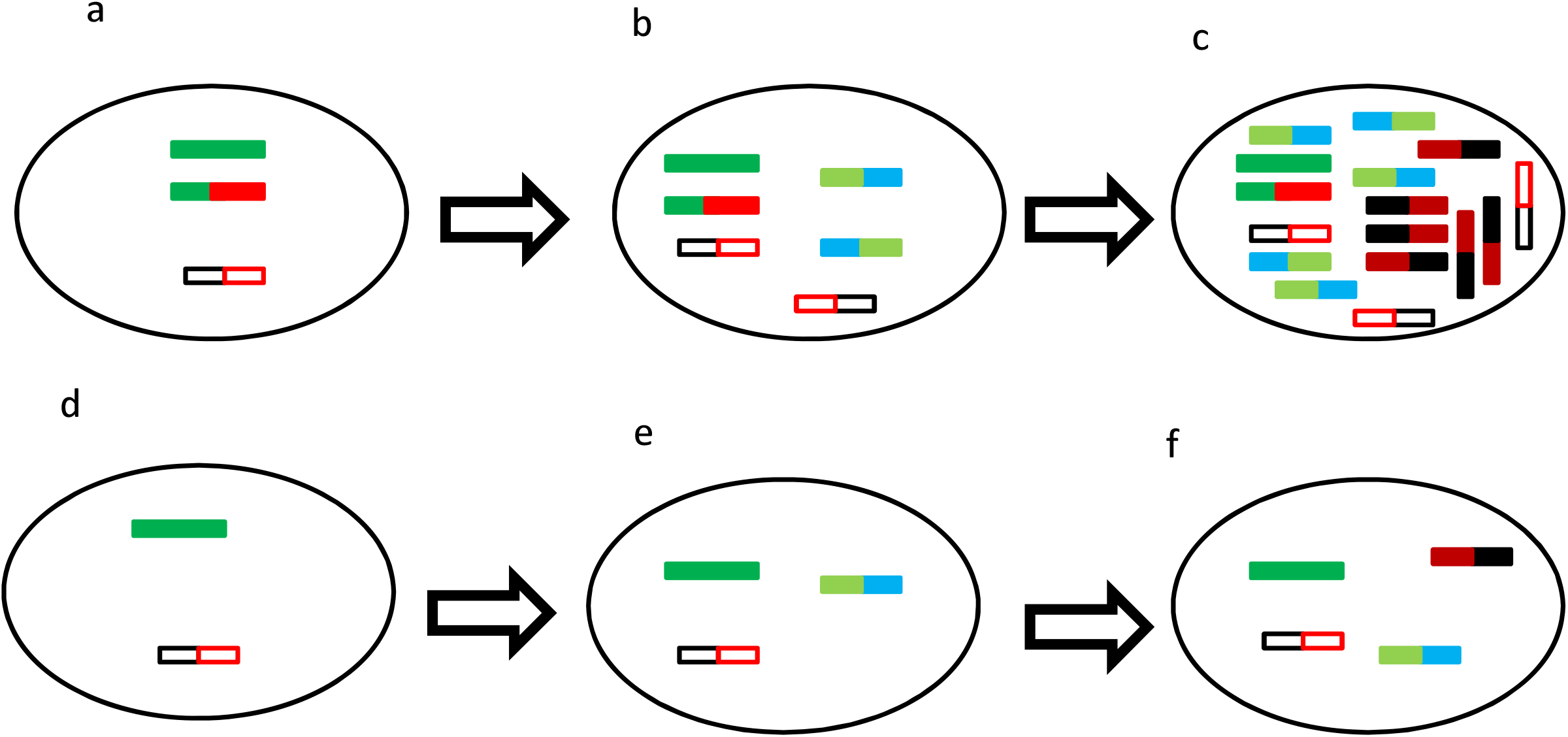
Interactions between HFGs and EFGs to initiate MM. a). Cells with *TPM4*-*KLF2* and other HFGs produced fusion or truncated proteins and altered gene expression; b). During their development, the cells with altered gene expression promoted EFG expressions such as *MTG1*-*SCART1* and *SCART1-CYP2E1* and maybe activated some dormant HFGs to be expressed; c). The cells with excessive EFG expressions resulted in genomic rearrangements producing somatic fusion genes. The cells began to accumulate somatic fusion genes and gene mutations and underwent natural selection. Finally, the cells were amplified uncontrollably and led to develop MM; d). The cells from healthy individuals had no key HFGs associated with MM, resulting in fewer gene expression errors; e). The cells with more minor expression errors resulted in fewer cellular restraints represented by fewer EFGs; f). The cells with fewer EFG expressions maintained normal DNA repair mechanisms and had fewer gene mutations and genomic rearrangements. The green rectangle indicates the normal genes. The solid red and green rectangle presented the key HFG, such as *TPM4*-*KLF2*. The open black and red box showed the minor sets of HFG. The light green and blue rectangles and blue and green rectangles represented EFGs such as *MTG1*-*SCART1* and *SCART1-CYP2E1*. The black and red rectangles and red and black rectangles indicated somatic fusion genes and gene mutations.

## Acknowledgments

We have expressed our most profound appreciation to Ms. Kathy Giust for her private information and support. We profoundly appreciate Prof. Yinxiong Li and his lab members’ contribution to this work.

## Figure Legends

**Supplementary Figure 1 Expression Levels of fusion transcripts with recurrent frequencies of ≥50%**. *KANSARL* isoforms 1 and 2 were used as references.

**Supplementary Figure 2 Hybrid protein encoded by *TPM4-KLF2* HFG** Underlined sequences represented the tropomyosin 4 amino acids. Asterisk was a stop codon.

Supplemental Table 1 Lists of fusion transcripts with recurrent frequencies of ≥10% discovered in 727 MMRFCS patients

Supplemental Table 2 Hereditary fusion gene list identified by overlapping between 1180 monozygotic twin hereditary fusion genes and fusion genes with recurrent frequencies of ≥10

Supplemental Table 3 Comparison of *MTG1-SCART1* and *SCART1-CYP2E1* recurrent frequencies between multiple myeloma and monozygotic twins

Supplemental Table 4 *MTG1-SCART1* Isoform lists discovered in 727 MMRFCS patients.

Supplemental Table 5 SCART1-CYP2E1 Isoform lists discovered in 727 MMRFCS patients.

## References

1. Raab MS, Podar K, Breitkreutz I, Richardson PG, Anderson KC. Multiple myeloma. Lancet. 2009;374(9686):324–339.

2. Mortality GBD, Causes of Death C. Global, regional, and national age-sex specific all-cause and cause-specific mortality for 240 causes of death, 1990-2013: a systematic analysis for the Global Burden of Disease Study 2013. Lancet. 2015;385(9963):117–171.

3. Caffrey MK. MMRF efforts attack the disease from every angle; a transformation is on the horizon. Am J Manag Care. 2014;20(11 Spec No.):E6.

4. Chapman MA, Lawrence MS, Keats JJ, et al. Initial genome sequencing and analysis of multiple myeloma. Nature. 2011;471(7339):467–472.

5. Egan JB, Shi CX, Tembe W, et al. Whole-genome sequencing of multiple myeloma from diagnosis to plasma cell leukemia reveals genomic initiating events, evolution, and clonal tides. Blood. 2012;120(5):1060–1066.

6. Lohr JG, Stojanov P, Carter SL, et al. Widespread genetic heterogeneity in multiple myeloma: implications for targeted therapy. Cancer Cell. 2014;25(1):91–101.

7. Rashid NU, Sperling AS, Bolli N, et al. Differential and limited expression of mutant alleles in multiple myeloma. Blood. 2014;124(20):3110–3117.

8. Tian E, Sawyer JR, Heuck CJ, et al. In multiple myeloma, 14q32 translocations are nonrandom chromosomal fusions driving high expression levels of the respective partner genes. Genes Chromosomes Cancer. 2014;53(7):549–557.

9. Manojlovic Z, Christofferson A, Liang WS, et al. Comprehensive molecular profiling of 718 Multiple Myelomas reveals significant differences in mutation frequencies between African and European descent cases. PLoS Genet. 2017;13(11):e1007087.

10. Cleynen A, Szalat R, Kemal Samur M, et al. Expressed fusion gene landscape and its impact in multiple myeloma. Nat Commun. 2017;8(1):1893.

11. Nasser S, Christofferson A, Legendre C, et al. Comprehensive Identification of Fusion Transcripts in Multiple Myeloma: An Mmrf Commpass Analysis. Blood. 2017;130:61.

12. Wang Y, Wu N, Liu J, Wu Z, Dong D. FusionCancer: a database of cancer fusion genes derived from RNA-seq data. Diagn Pathol. 2015;10:131.

13. Prideaux SM, Conway O’Brien E, Chevassut TJ. The genetic architecture of multiple myeloma. Adv Hematol. 2014;2014:864058.

14. Mertens F, Johansson B, Fioretos T, Mitelman F. The emerging complexity of gene fusions in cancer. Nat Rev Cancer. 2015;15(6):371–381.

15. Ge H, Liu K, Juan T, Fang F, Newman M, Hoeck W. FusionMap: detecting fusion genes from next-generation sequencing data at base-pair resolution. Bioinformatics. 2011;27(14):1922–1928.

16. Haas BJ, Dobin A, Li B, Stransky N, Pochet N, Regev A. Accuracy assessment of fusion transcript detection via read-mapping and de novo fusion transcript assembly-based methods. Genome Biol. 2019;20(1):213.

17. Hu X, Wang Q, Tang M, et al. TumorFusions: an integrative resource for cancer-associated transcript fusions. Nucleic Acids Res. 2018;46(D1):D1144–D1149.

18. Zhou JX, Yang X, Ning S, et al. Identification of KANSARL as the first cancer predisposition fusion gene specific to the population of European ancestry origin. Oncotarget. 2017;8(31):50594–50607.

19. Qin F, Song Z, Babiceanu M, et al. Discovery of CTCF-sensitive Cis-spliced fusion RNAs between adjacent genes in human prostate cells. PLoS Genet. 2015;11(2):e1005001.

20. Grosso AR, Leite AP, Carvalho S, et al. Pervasive transcription read-through promotes aberrant expression of oncogenes and RNA chimeras in renal carcinoma. Elife. 2015;4.

21. Thierry-Mieg D, Thierry-Mieg J. AceView: a comprehensive cDNA-supported gene and transcripts annotation. Genome Biol. 2006;7 Suppl 1:S12 11–14.

22. Zhuo D. The First Glimpse of Homo sapiens Hereditary Fusion Genes. Journal of Pharmacogenomics & Pharmacoproteomics. 2022;13(4):1–7.

23. Mitelman F, Johansson B, Mertens F, Schyman T, Mandahl N. Cancer chromosome breakpoints cluster in gene-rich genomic regions. Genes Chromosomes Cancer. 2019;58(3):149–154.

24. Nakai K, Horton P. PSORT: a program for detecting sorting signals in proteins and predicting their subcellular localization. Trends Biochem Sci. 1999;24(1):34–36.

25. Roberts KG, Morin RD, Zhang J, et al. Genetic alterations activating kinase and cytokine receptor signaling in high-risk acute lymphoblastic leukemia. Cancer Cell. 2012;22(2):153–166.

26. Irene J. Locher WA, Daniel M. Borràs, M. Willy Honders, Rick H. de Leeuw, Wilma G.M. Kroes, Peter de Knijff, Cornelis A.M. van Bergen, Peter A.C. ‘t Hoen, Szymon M. Kielbasa, Jeroen F.J. Laros, Marieke Griffioen, Hendrik Veelken,. Fusion Transcripts without Corresponding Cytogenetic Abnormalities in Acute Myeloid Leukemia: Implications for AML Pathogenesis,. Blood. 2017;130(Supplement 1,):2703.

27. Tajiri R, Shiba E, Iwamura R, et al. Potential pathogenetic link between angiomyofibroblastoma and superficial myofibroblastoma in the female lower genital tract based on a novel MTG1-CYP2E1 fusion. Mod Pathol. 2021;34(12):2222–2228.

28. Kuhn J, Meerzaman D, Ries RE, et al. PIM3-SCO2 Fusion Is a Novel Transcription-Induced Chimera That Is Highly Prevalent In Childhood AML. Blood. 2013;122(21):2549–2549.

