## Supplementary for "The Hereditary and Epigenetic Fusion Gene Signatures of Multiple Myeloma"

Supplementary Figure 1

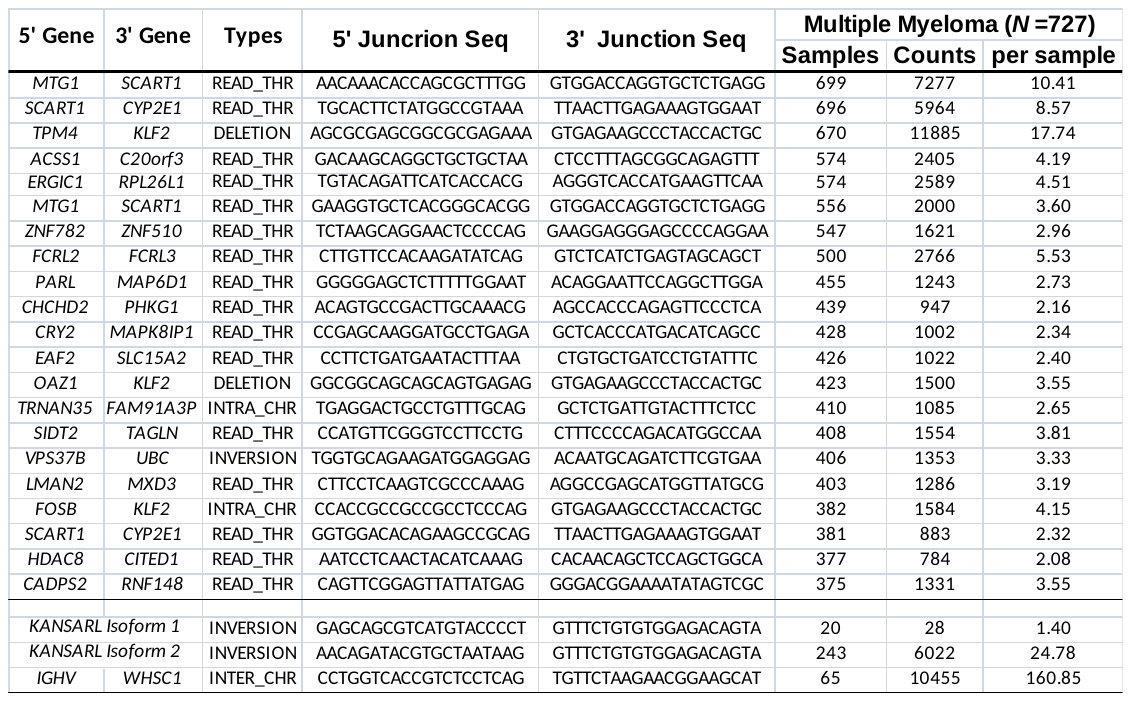

**Supplementary Figure 1 Expression Levels of fusion transcripts with recurrent frequencies of ≥50%.** *KANSARL* isoform 1 and 2 were used references.

Supplementary Figure 2

MAGLNSLEAVKRKIQALQQQADEAEDRAQGLQRELDGERERREKVRSPTTATGTAAAGSLRAQTSSRATTESTRATGHSSAICAIVPSRAPITWRCT*

**Supplementary Figure 2 Hybrid protein encoded by *TPM4-KLF2*** **HFG** Underlined sequences represented the tropomyosin 4 amino acids. Asterisk was a stop condon.
